# Growth hormone receptor (GHR)-expressing neurons in the hypothalamic arcuate nucleus regulate glucose metabolism and energy homeostasis

**DOI:** 10.1101/2020.08.17.254862

**Authors:** Juliana Bezerra Medeiros de Lima, Lucas Kniess Debarba, Manal Khan, Chidera Ubah, Olesya Didyuk, Iven Ayyar, Madelynn Koch, Marianna Sadagurski

## Abstract

Growth hormone (GH) receptor (GHR), expressed in different brain regions, is known to participate in the regulation of whole-body energy homeostasis and glucose metabolism. However, GH activation of these GHR-expressing neurons is less studied. We have generated a novel GHR-driven Cre recombinase transgenic mouse line (GHR^cre^) in combination with the floxed tdTomato reporter mouse line we tracked and activated GHR-expressing neurons in different regions of the brain. We focused on neurons of the hypothalamic arcuate nucleus (ARC) where GHR was shown to elicit a negative feedback loop that regulates GH production. We found that ARC^GHR+^ neurons are co-localized with AgRP, GHRH, and somatostatin neurons, which were activated by GH stimulation. Using designer receptors exclusively activated by designer drugs (DREADDs) to control GHR^ARC^ neuronal activity, we revealed that activation of GHR^ARC^ neurons was sufficient in regulating distinct aspects of energy balance and glucose metabolism. Overall, our study provides a novel mouse model to study *in vivo* regulation and physiological function of GHR-expressing neurons in various brain regions. Furthermore, we identified for the first time specific neuronal population that responds to GH and directly linked it to metabolic responses *in vivo*.

## Introduction

A cumulative body of evidence established that growth hormone (GH) plays pivotal roles in the regulation of systemic metabolism, through activation of the GH receptor (GHR) in the liver, muscle, adipose, and other tissues (1-6). In the central nervous system (CNS), GH is present in regions known to participate in the regulation of feeding, energy balance, and glucose metabolism, including the hypothalamus, hippocampus, and amygdala (7-11). The expression of GHR within the CNS has been mapped by *in situ* hybridization and by detection of the downstream target, the phosphorylated activator of transcription (STAT) 5, revealing large numbers of GH-responsive neurons in various brain regions (8,12,13). While these studies detected GHR expression within the CNS, the functional assessment of the GHR-expressing neurons in various brain regions was lacking. GHR expression in the brain is critical for the neuroendocrine neurons to sense and regulate GH production by the pituitary (8,14,15). In the arcuate nucleus of the hypothalamus (ARC), the GHR is involved in a negative feedback loop that regulates GH production and secretion by GH-releasing hormone (GHRH) (8). As part of this negative feedback, GH inhibits it’s own secretion acting on the GHR in neuropeptide Y (NPY) neurons in the ARC and somatostatin (SST) neurons in the paraventricular nucleus (PVN). Activation of these neurons augments SST release and inhibits GH secretion (16,17). In recent years, it became clear that GH action in the ARC represents an important component of energy homeostasis (13). We have recently shown that neuronal-specific deletion of GHR in leptin receptor (LepRb)-expressing neurons in the hypothalamus impaired hepatic glucose production and systemic lipid metabolism (18). Additionally, mice lacking GHR specifically in the orexigenic agouti-related peptide (AgRP) expressing neurons in the ARC display impaired responses to fasting and food restriction, while deletion of GHR from anorexigenic proopiomelanocortin (POMC) neurons in the ARC did not produce significant metabolic phenotype (19,20). Collectively, these results indicated unique roles of GHR-expressing neurons in the ARC in metabolic control. However, in vivo GH-mediated activation of GHR-expressing neurons were not studied.

In the current study, we specifically aimed at studying in vivo activation of these complex neural circuitry of the GHR-expressing neurons in the ARC. To that end, we developed a novel GHR-driven cre mouse (GHR^cre^) using the CRISPR/Cas9 gene-editing technology. The new GHR^cre^ model allowed us to both track and activate GHR-expressing neurons. Utilizing these mice, we studied the functional roles of GHR neurons in the ARC in the regulation of systemic glucose metabolism and energy homeostasis. We found that activation of GHR^ARC^ neurons acutely increased systemic glucose sensitivity, energy expenditure, and heat production. Overall, our study revealed a novel network of metabolic regulation through the hypothalamic GH axis in the ARC. Finally, our mouse model provides a novel tool to identify specific neuronal populations mediating the effects of GH in different brain regions.

## Materials and Methods

### Experimental Animals

GHR^cre^ mice were generated using the Clustered Regularly Interspaced Short Palindromic Repeats associated protein Cas9 (CRISPR/Cas9) technology (21,22). All procedures were performed at the University of Michigan Transgenic Core as before (22). A detailed description of the procedures is described in Supplementary Materials and Methods. tdTomato mice on the ROSA26 background (B6.Cg-Gt(ROSA)26Sort^m14(CAG-tdTomato)Hze^/J, (stock 007914) were purchased from The Jackson Laboratory. Adult male mice (8-12 weeks old) were used for all studies. All mice were provided *1a9T d libitum19T* access to standard chow diet (Purina Lab Diet 5001) and housed in temperature-controlled rooms on a 12-hour/12-hour light-dark cycle. Mice were bred and housed within our colony according to guidelines approved by the Wayne State University Committee on the Care and Use of Animals.

### Perfusion and Histology

Mice were anesthetized (IP) with avertin and transcardially perfused with phosphate-buffered saline (PBS) (pH 7.5) followed by 4% paraformaldehyde (PFA). Brains were post-fixed, sank in 30% sucrose, frozen in OCT medium, and then sectioned coronally (30 µm) and processed for immunohistochemistry as previously described (23,24). For immunohistochemistry, free-floating brain sections were washed in PBS, blocked using 3% normal donkey serum (NDS) and 0.3% Triton X-100 in PBS and then stained with a primary antibody for 48 hours at 4°C with agitation in blocking buffer: DsRed (anti-rabbit, 1:5000, cat. number NC9580775, Takara), GFP (anti-chicken, 1:1000, cat. number ab13970, Abcam), anti-tdTom (anti-goat, 1:500, cat. number AB8181-200, Scigen), pSTAT5 (anti-rabbit, 1:500, cat. number 9359, Cell Signaling), GFAP (anti-chicken, 1:500, cat. number Ab5541, Millipore), Iba-1 (anti-goat, 1:1000, cat. number ab5076, Abcam) and cFos (anti-sheep, 1:500, cat. number ab6167, Abcam) For pSTAT5 staining sections were pretreated for 10 min in 90% Methanol and 10% H_2_O_2_ in PBS before blocking buffer incubation. On the following day, all floating brain sections were washed with PBS 0.1M and incubated with the following secondary antibodies for 2 hours: donkey anti-rabbit, anti-goat, anti-sheep, anti-chicken Alexa Fluor 488 and/or 568 (Invitrogen, 1:200). For the staining specificity control, the immunohistochemical experiments were performed with brain sections in which the primary antibody was omitted and substituted with serum.

### Two-plex fluorescent in-situ hybridization

Fixed-frozen ARC-containing GHR^cre^ brain sections of 12-week old male mice (10µm) were processed for the RNAscope Fluorescent Multiplex assay (Advanced Cell Diagnostics, Inc). The samples were double-labeled with probes for GHR (Mm-Ghr-C2 464951), GHRH (Mm-Ghrh-C2 470991), or SST (Mm-Sst-C2 404631) together with tdTom (tdTomato-C3 317041).

### Images and data analysis

All sections used for ISH were visualized with a Zeiss M2 microscope blindly. All other fluorescent sections were visualized with a Nikon Eclipse Ni microscope coupled to a Nikon DS-Ri2 camera. Photomicrographs were captured using the NIS-Elements Br 5.0 Zen software. Fiji ImageJ image-editing software was used to overlay photomicrographs to construct merged images and to mount plates. Only sharpness, contrast, and brightness were adjusted and the same values for each target labeled were applied.

### Surgery and viral injections

Stereotaxic viral injections were performed as described (25). Briefly, animals were anesthetized using 1-3% isoflurane, their head shaved and placed in a three-dimensional stereotaxic frame (Kopf 1900, Cartesian Research Inc., CA). The skull was exposed with a small incision, and two small holes were drilled for bilateral microinjection (200 nL/side) of the excitatory DREADD, AAV8-hSyn-DIO-hM3DGq-mCherry (cat. number # 44361-AAV8, Addgene) into the ARC of GHR^cre^ mice at stereotaxic coordinates based on the Mouse Brain Atlas: A/P: −1.3, M/L: +/-0.2, D/V: −5.85 (26). Animals received a pre-operative dose of buprenorphine hydrochloride (1 mg/kg). After surgery, mice were allowed 2 weeks of recovery to maximize virally-transduced gene expression and to acclimate animals to handling and experimental paradigms before the study. Activation of the DREADD receptor was induced by intraperitoneal administration of the agonist, clozapine-N-oxide (CNO, 0.3 mg/kg, ip, cat. number 4936, Tocris). Expression was verified post hoc in all animals, and any data from animals in which the transgene expression was located outside the targeted area were excluded from analysis.

### Metabolic Analysis

Following recovery, GHR^cre^ mice with activating DREADD (hM3Dq) underwent glucose metabolism and energy expenditure assays as previously described (27). Intraperitoneal glucose tolerance tests were performed on mice fasted for 6 hours. Mice were administered with 0.9% saline or CNO 1 hour before glucose (2 g/kg BW) injection. GTTs tests were performed one week apart and blood glucose levels were measured as before (28). Blood insulin was determined using a Mouse Insulin ELISA kit (cat. number 50-194-7920, Crystal Chem. Inc.). For peripheral GH stimulation (recombinant mouse GH, 12.5 µg/100g BW, National Hormone & Peptide Program, Harbor-UCLA Medical Center, CA), mice were injected i.p. and perfused 1.5 hours later, for pSTAT5 immunostaining as described before (18). Metabolic measurements of energy homeostasis were obtained using an indirect calorimetry system (PhenoMaster, TSE system, Bad Homburg, Germany). The mice were acclimatized to the cages for 3 days and monitored for 5 days, food and water were provided *ad libitum*. Following acclimatization, GHR^cre^ excitatory DREADD-expressing mice received an i.p. injection of vehicle (0.9% saline) and measurements were analyzed for the following 8 hours. Mice remained in the chambers with food and water *ad libitum* and 72 hours later the same experimental design was repeated, and animals were treated with an i.p. injection of CNO (0.3 mg/kg). Data were analyzed vehicle vs CNO per mouse.

### Statistical analysis

Unless otherwise stated mean values ± SEM is presented in graphics. GTT data were analyzed by residual maximum likelihood (REML) mixed model followed by Sidak’s post hoc while cumulative RER, heat production, ambulatory activity, food intake, and water intake data were analyzed through paired t-test. Post-hoc comparisons were only carried out when the p-value was significant for effect and/or interactions. *p* < 0.05 was considered statistically significant.

## Results

### Characterization of the GHR^cre^ mice

To characterize the role of GHR-expressing neurons in the ARC, we have developed a GHR cell-specific molecular tool (GHR^cre^) using CRISPR/Cas9 gene-editing technology (Supplementary Figure 1A). The GHR^cre^ mice reproduced in Mendelian ratio and both male and female mice exhibited normal body weight, and fed/fasting blood glucose levels (Supplementary Figure 1B and 2). GHR^cre^ mouse line was validated by a cre-dependent Rosa26-tdTomato reporter mouse. The expression pattern of tdTomato reporter revealed the presence of GHR^cre^-expressing neurons in the several areas of the hypothalamus (Figure 1A), including the midbrain and hindbrain (Supplementary Figure 1C and Supplementary Table 1 and 2) (8,13). To validate the expression of GHR in our GHR^tdTom^ mice, we performed RNA *in situ* hybridization with RNAscope, using probes against *GHR* and *tdTomato* in the hypothalamic arcuate nucleus (ARC). As seen in Figure 1B, the majority of TdTomato^+^ neurons were positive for the expression of the *GHR* gene.

**Table 1.**
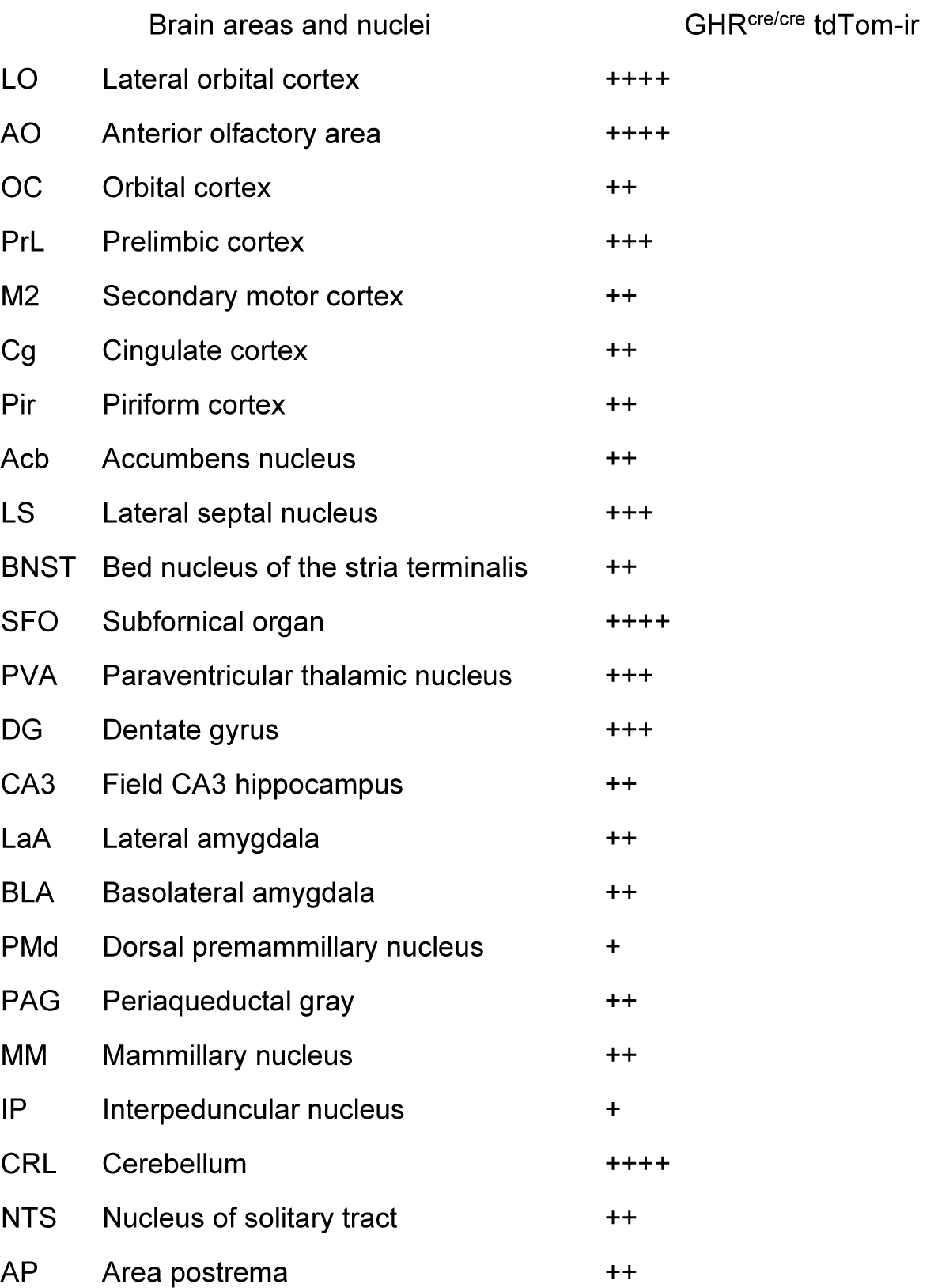
Distribution of GHR^cre/cre^ TDTom immunoreactive neurons

**Table 2.** Distribution of GHR^cre/cre^ TDTom immunoreactive neurons

Distribution of GHR<sup>cre/cre</sup> TDTom immunoreactive neurons
|  | hypothalamic areas and nuclei | GHR<sup>cre/cre</sup> tdTom-ir |
| --- | --- | --- |
| MPOa | Medial preoptic area | + |
| RCH | Retrochiasmatic area | ++ |
| PVH | Paraventricular nucleus | ++ |
| ARH | Arcuate nucleus | ++ |
| VMH | Ventromedial nucleus | +++ |
| DMh | Dorsomedial nucleus | +++ |
| LHA | Lateral hypothalamic area | + |
| DA | Dorsal hypothalamic area | + |
| pHA | Posterior hypothalamic area | +++ |
| pmV | Ventral premammillary nucleus | ++ |

**Figure 1.**
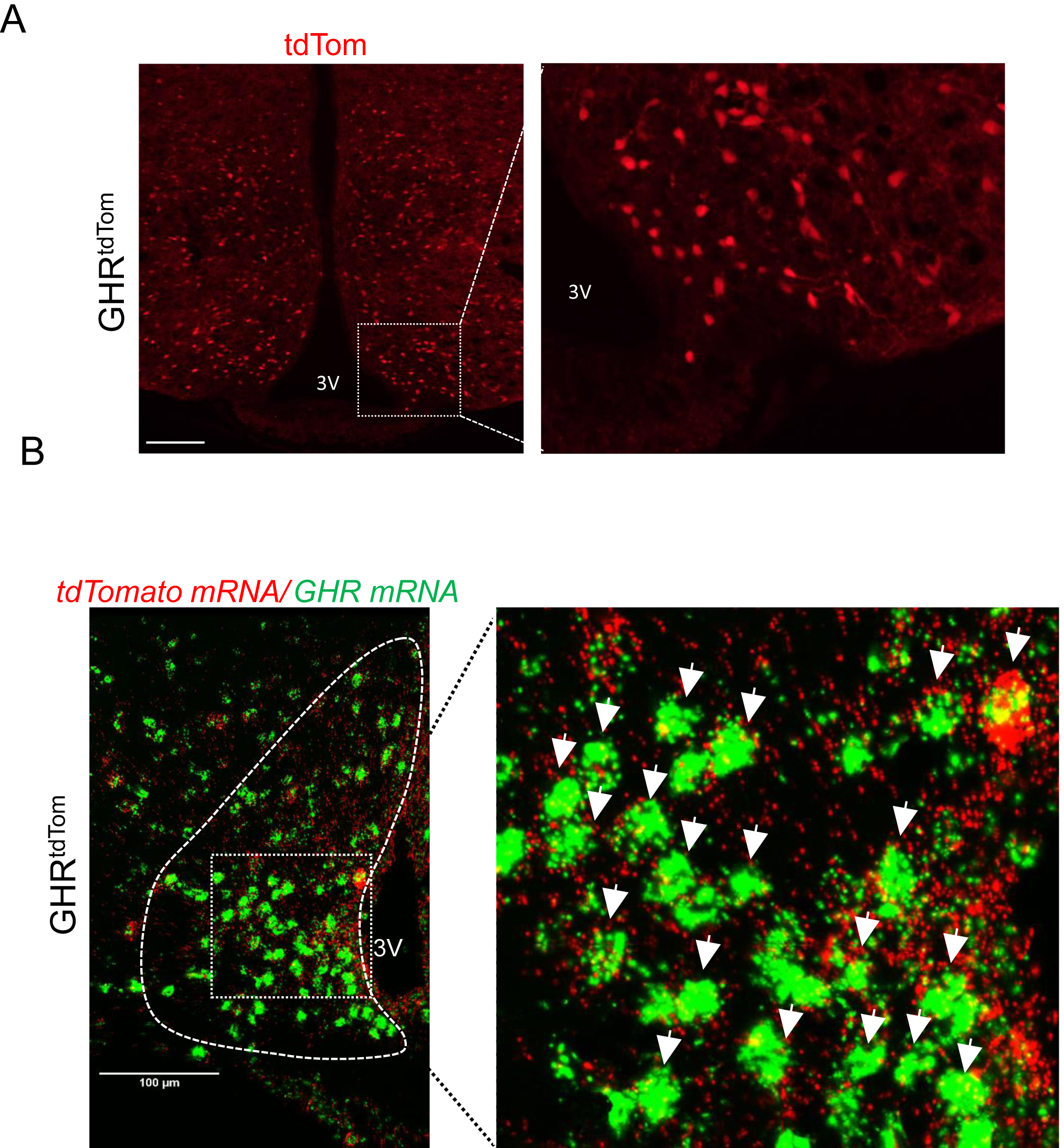
GHR-expressing neurons in the hypothalamus. To visualize cre-expressing neurons, mice were crossed with tdTomato reporter mice. (A) Immunofluorescent image of GHR^+^ neurons in the hypothalamus (red, TdTomato). The dashed box indicates a region of the arcuate nucleus of the hypothalamus (ARC) that is digitally enlarged and shown as an inset. (B) Two-plex fluorescent *in situ* hybridization of *GHR* mRNA (green) and *tdTomato* mRNA (red) was performed on coronal slices in the ARC. The dashed box indicates the region of the ARC that is digitally enlarged and shown as inset demonstrating the colocalization of *GHR* and *tdTomato* mRNA (white arrows). 3V = Third ventricle. Scale bar: 100 µm.

Upon binding to the GHR, GH triggers the activation of the JAK/Stat5 pathway (4,29,30). To track GH-mediated STAT5 phosphorylation (pSTAT5), acute intraperitoneal GH injection was given to the GHR^tdTom^ mice. We found that pStat5 was colocalized with the majority of the ARC^GHR+^ neurons (Figure 2A). We did not detect the colocalization of GHR^tdTom+^ cells in astrocytes positive to glial fibrillary acidic protein (GFAP). Additionally, the Iba1, a marker of microglia, did not co-localize with tdTomato (Figure 2B), indicating that GHR signaling is principally targeting neurons and not glia cells.

**Figure 2.**
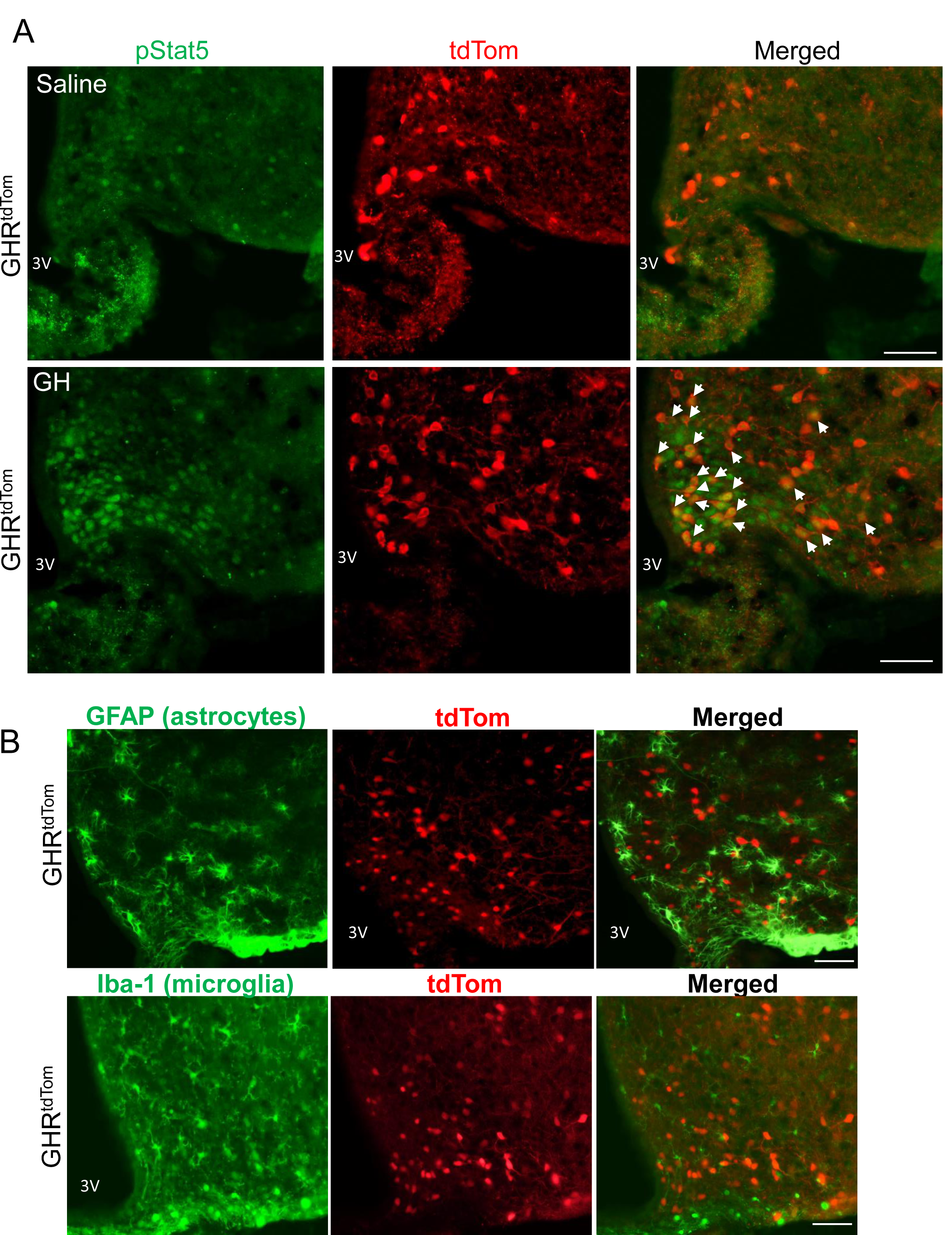
Characterization of GHR-expressing neurons in the ARC. (A) GH signaling in GHR-expressing neurons in the ARC. Immunofluorescence for pSTAT5 in 12-week-old GHR^tdTom^ mice injected IP with vehicle (saline) or GH (12.5µg/100g BW; 1.5 hr). Representative images from the ARC of GHR^tdTom^ mice are shown. pSTAT5 (green), TdTomato (red), and merged images of the indicated mice (colocalization is shown by arrows). (B) Representative images of astrocytes identified by immunofluorescent detection of GFAP protein (green, upper panel) and microglia evaluated by Iba1 immunostaining (green, lower panel) in the ARC obtained from GHR^tdTom^ mice (red, TdTomato). 3V = Third ventricle. Scale bar: 100 µm.

To determine whether GHR^ARC+^ neurons overlap with other known ARC populations that are involved in neuroendocrine regulation, we further examined the expression of *SST* and *GHRH* in identified GHR^ARC+^ neurons using double IHC and ISH. We found two-plex fluorescent ISH for *GHRH* or *SST* and *tdTomato* mRNAs colocalized in the ARC of GHR^tdTom^ mice (Figure 3), as previously reported (12). In support of previous studies (19,31), we further confirmed a substantial overlap of GHR^+^ neurons in the ARC with AgRP-expressing neurons in the GHR^tdTom^ *P*mice (Supplementary Figure 3A). Additionally, in support of the single-cell sequencing data (31), we found a minimal colocalization with dopaminergic neurons (GHR^tdTom+^/TH^+^ cells), and with POMC neurons (GHR^tdTom+^/β-endorphin^+^ cells) in the ARC (Supplementary Figure 3B and 3C).

**Figure 3.**
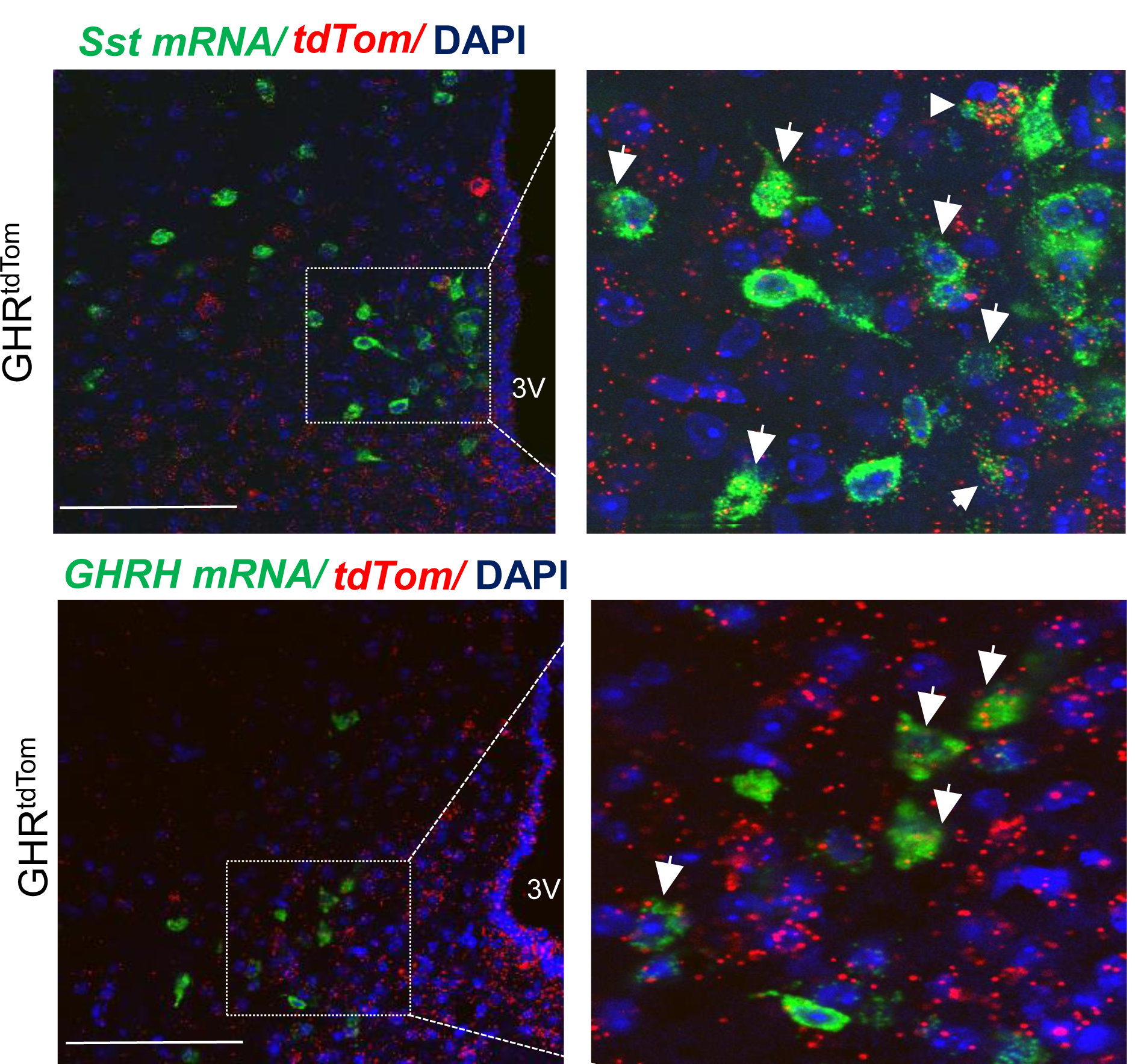
GHR-expressing neurons colocalization with SST and GHRH in the ARC. Two-plex fluorescent *in situ* hybridization of (A) *SST* mRNA (green), *tdTomato* mRNA (red) and *DAPI* (blue), and (B) *GHRH* mRNA (green), *tdTomato* mRNA (red) and *DAPI* (blue) was performed on coronal slices containing the ARC. Dashed box indicates the region of the ARC that is digitally enlarged and shown as inset demonstrating the colocalization of *SST* and *tdTomato* mRNA or *GHRH* and *tdTomato*. 3V = Third ventricle. Scale bar: 100 µm.

### GHR^ARC+^ neurons regulate glucose metabolism

The hypothalamic GHR-expressing neuronal circuits operate within complex physiological settings involving the interrelationships between SST, GHRH, and AgRP-expressing neurons. We aimed to directly assess the contribution of ARC GHR-expressing neuronal populations to glucose metabolism and energy homeostasis. To achieve that, we employed a Cre-dependent DREADD (Designer Receptors Exclusively Activated by Designer Drugs) virus to acutely modulate neuronal activity in response to peripheral injection of an otherwise inert compound, clozapine N-oxide (CNO) (32-34). To determine whether activation of GHR^ARC+^ neurons can influence blood glucose levels, we enhanced the GHR^ARC+^ neuronal activity of GHR^Cre^ mice by stereotactically injecting AAV8-DIO-hM3Dq-mCherry into the ARC and activated the transduced cells with CNO (Figure 4A). Specific activation of GHR^ARC^ neurons was demonstrated by nuclear c-Fos expression as a marker of neuronal activation in AAV8-DIO-hM3Dq-mCherry-ARC-injected GHR^cre^ mice treated with vehicle (data not shown) or CNO (Figure 4A) before perfusion. CNO administration resulted in a significant increase in c-Fos expression in hM3Dq-expressing ARC neurons. Basal blood glucose and serum insulin concentrations were indistinguishable between baseline and CNO injected mice (Figure 4B and D). Each animal served as its own control (e.g., saline versus CNO). Despite unchanged fasting blood glucose levels, AAV8-DIO-hM3Dq-mCherry-ARC CNO-treated GHR^cre^ mice displayed significantly increased glucose disposal, indicating increased sensitivity in response to an intraperitoneal glucose load (AUC baseline: 1234 ± 88. 67 vs AUC CNO: 1049 ± 84. 27, t-test *p* < 0.05, Figure 4B and C). Of note, the DREADD virus per se (off-target infection, with or without CNO) did not affect glucose tolerance (data not shown).

**Figure 4.**
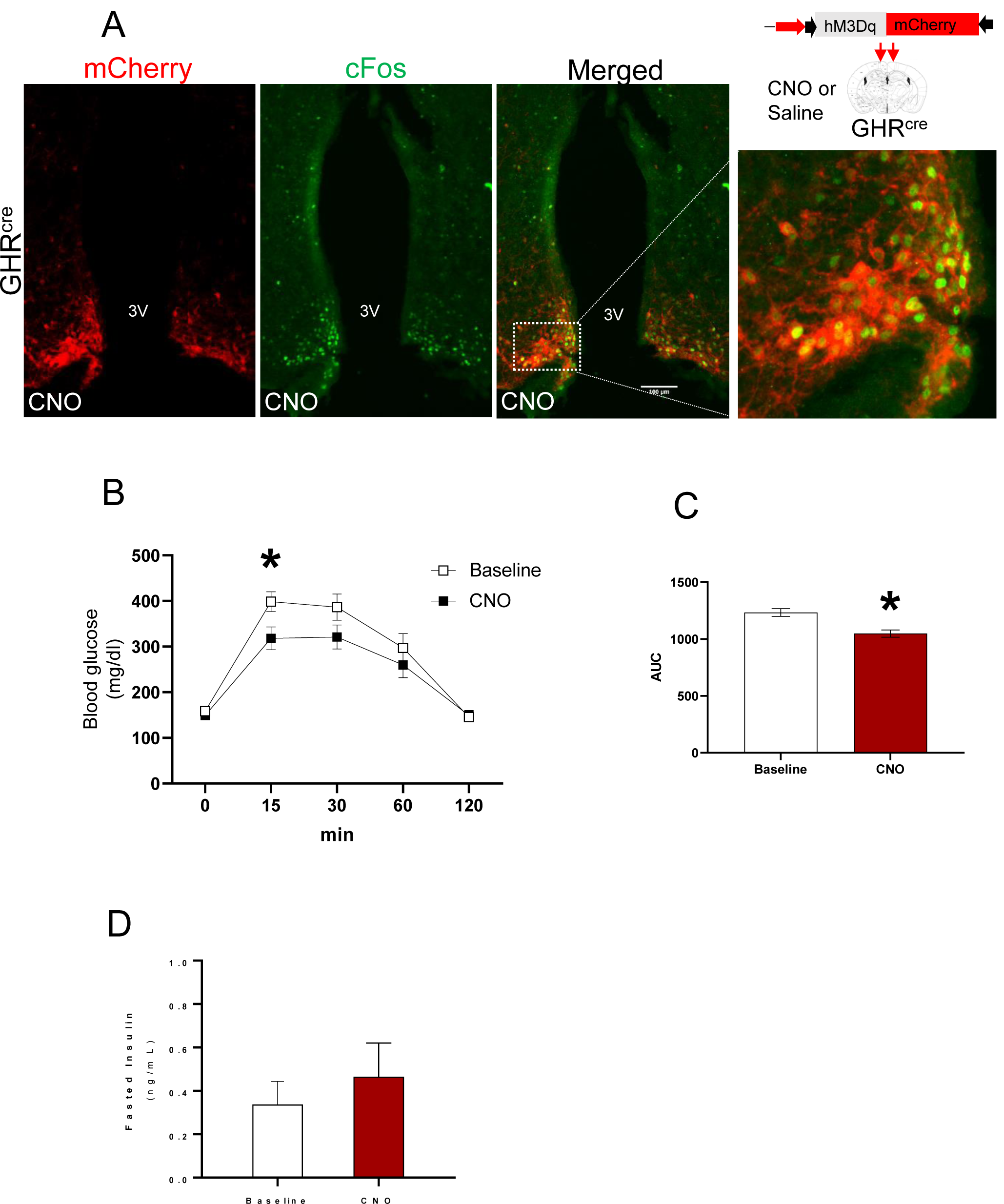
Acute activation of GHR^ARC^ neurons increases glucose tolerance. (A) Neuronal activation by cFos (green) was assessed 90 minutes after CNO stimulation. IHC for mCherry (red) identifies AAV-hM3Dq expression in GHR^ARC^ neurons in the ARC. Merged image (green/red) and dashed box indicate the region of the ARC that is digitally enlarged and shown as inset demonstrating the colocalization of cFos and mCherry. 3V =Third ventricle. Scale bar: 100 µm. (B) Glucose tolerance tests (GTT) of 12-week old male mice performed one week apart. Saline (0.1 mL/10 g BW – Baseline) or CNO (0.3 mg/kg BW i.p) was injected 1 hour before i.p. GTT. The effect of GHR^ARC^ activation was analyzed by residual maximum likelihood (REML) mixed model followed by Sidak’s post hoc. (C) GTT area under the curve (AUC) was analyzed by a paired t-test. (D) Fasted insulin levels. Results are presented as mean ± SEM, n=7; * *p* < 0.05 compared to vehicle values

### GHR^ARC+^ neurons regulate energy balance and heat production

To establish the significance of GHR^ARC+^ neurons in the control of energy utilization, we analyzed components of energy expenditure in *ad libitum*-fed 12-week-old male GHR^cre^ mice. We injected mice with CNO in the morning during the light cycle, a time in which mice normally refrain from eating. Using a single-subject approach where each mouse serves as its own control, we showed that stimulation of GHR^ARC+^ neurons produced a significant increase in energy expenditure (Figure 5A), which lasted for approximately 8 hours. This effect was also associated with a significant increase in heat production in these CNO stimulated hM3Dq-ARC-injected GHR^cre^ mice (paired t-test, *p* < 0.05) (Figure 5B). CNO stimulated locomotor activity was equivalent to the baseline measurements (Figure 5C). Notably, acute activation of AgRP neurons markedly reduces energy expenditure (25), emphasizing the complexity of ARC GH-responsive neurons, and suggesting that GHR^ARC+^/AgRP^-^ neurons are critical to driving energy homeostasis.

**Figure 5.**
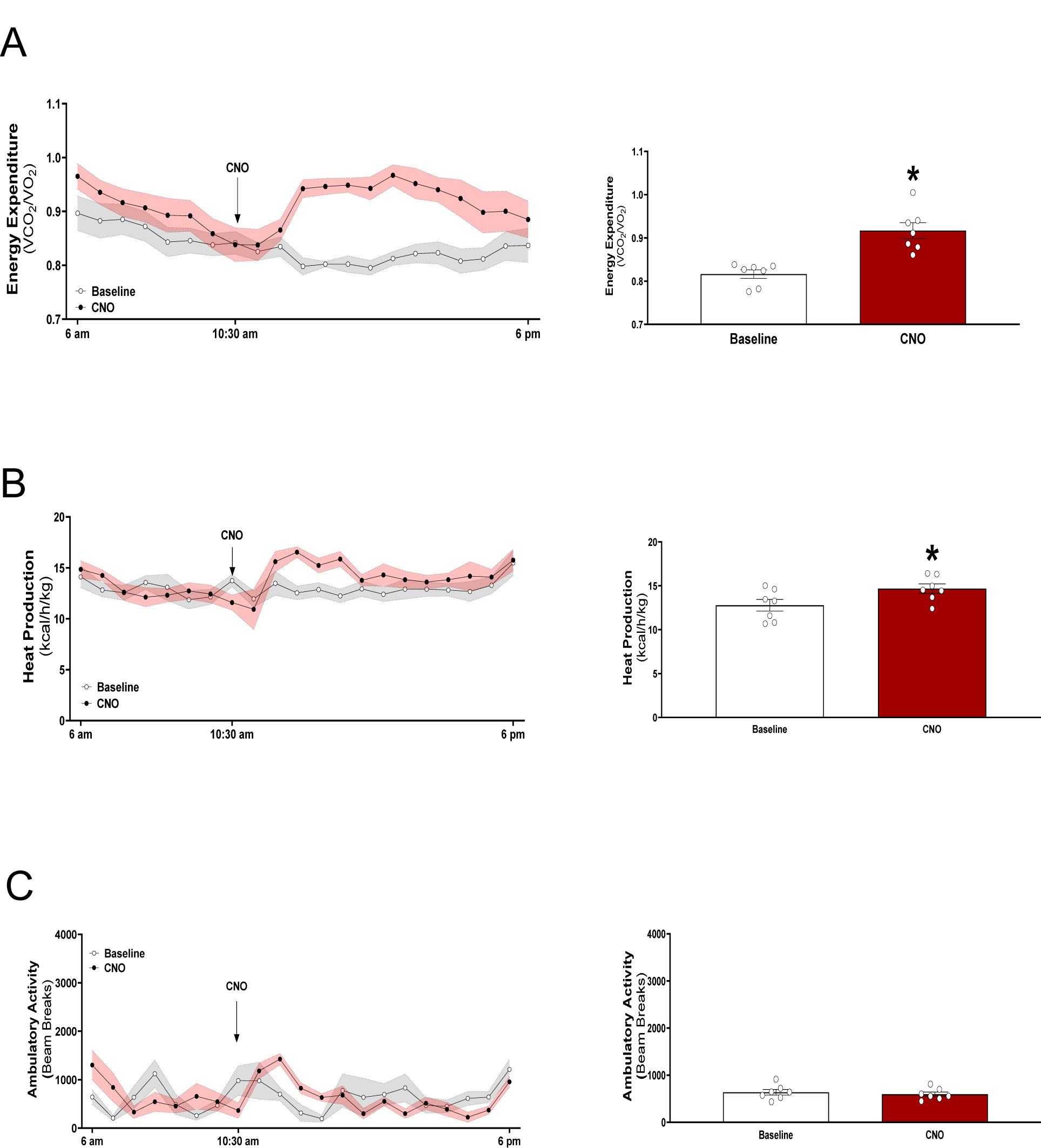
Acute activation of GHR^ARC^ neurons increases energy homeostasis but not ambulatory activity. (A) Respiratory exchange ratio (RER). (B) Heat production. (C) Ambulatory activity assessed by a total of beam breaks. Mice were acclimated in metabolic cages and i.p. injected with either saline (grey) or CNO (red) at 10:30 am. On the right side, AUC of the light cycle period from the treatment time. Data are from male mice, analyzed by paired t-test (mean ± SEM, n = 7; * *p* < 0.05 compared to vehicle values)

Surprisingly, administering CNO to these mice acutely and significantly increased feeding and drinking responses (Figure 6A and B), suggesting that GHR^ARC+^ neurons are orexigenic neurons functionally similar to AgRP/SST neuronal cluster in the ARC (31).

**Figure 6.**
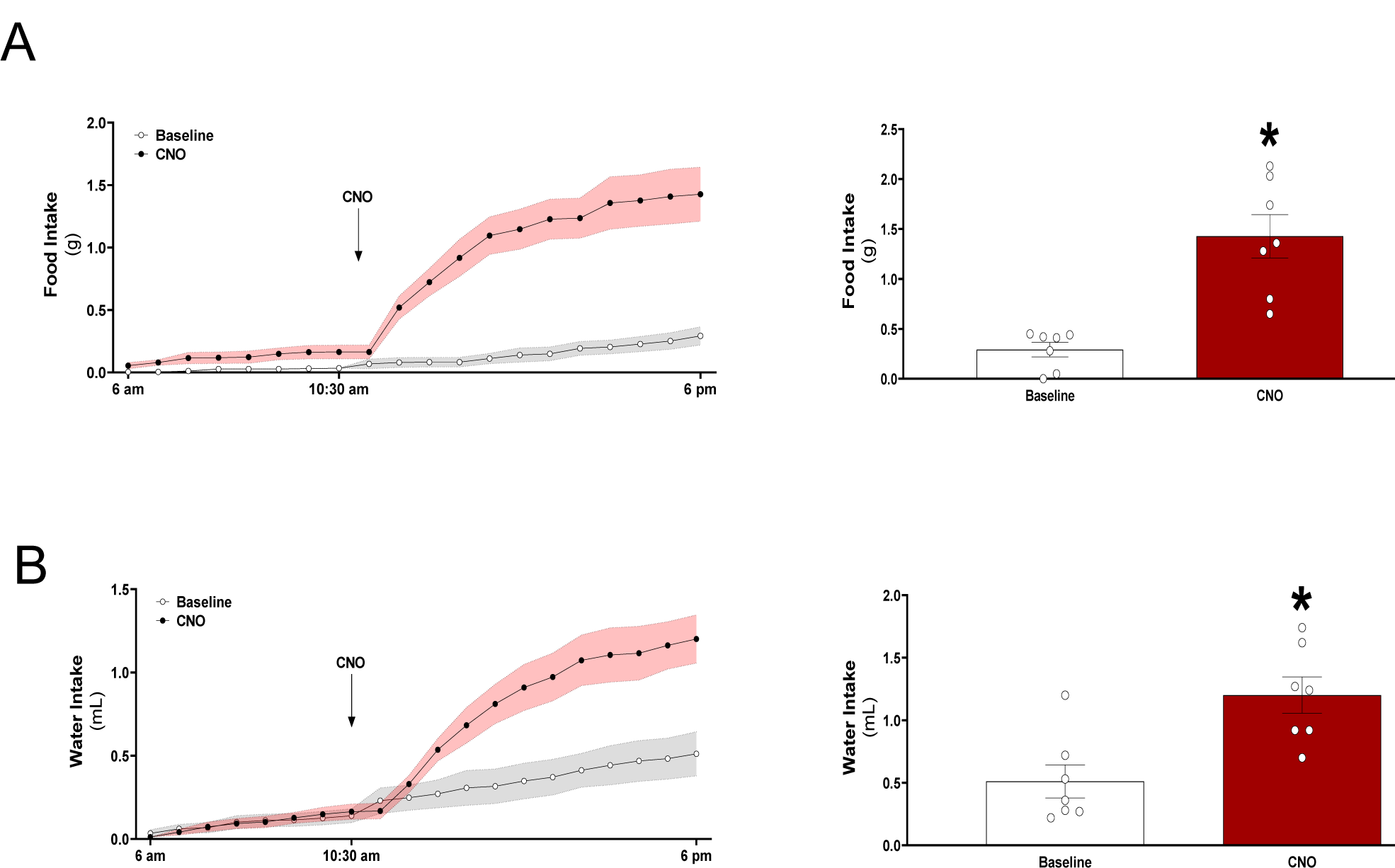
Acute activation of GHR^ARC^ neurons increases food and water intake. (A) Food intake. (B) Water intake. Mice were acclimated in metabolic cages and i.p. injected with either saline (grey) or CNO (red) at 10:30 am. Results are presented as mean ± SEM (filled area). On the right side, AUC of the light cycle period from the treatment time. Data are from male mice, analyzed by paired t-test (mean ± SEM, n = 7; * *p* < 0.05 compared to vehicle values

## Discussion

We present herein a novel mouse model that expresses cre recombinase driven by the GHR promoter. Specifically, using tdTomato immunoreactivity as a marker of GHR expression in GHR^tdTom^ mice, we unraveled the uncharacterized population of GHR expressing neurons in the ARC. Further, using a site-specificity approach we have characterized the function of GHR^+^ population in the ARC. Using a combination of a genetic mouse model with site-specific delivery of chemogenetic agent (CNO) we identified, for the first time, a GHR^ARC+^ neuronal population that plays a critical role in the maintenance of peripheral glucose metabolism and energy homeostasis.

The distribution of GHR-expressing neurons in the brain by GHR^dTom^ reporter resembles that determined by *in situ* hybridization and by systemic GH injections followed by pSTAT5 expression pattern in the brain (8,12,13). Large populations of GHR-expressing neurons lay in the hypothalamus, especially in the ARC, DMH, and VMH; other substantial populations reside in the posterior hypothalamic area and ventral pre-mammillary nucleus. The hippocampus areas, the cortex, the cerebellum, and the olfactory area also contain substantial concentrations of GHR-expressing neurons. The nucleus tractus solitarius (NTS) represents the hindbrain site with significant numbers of GHR-expressing neurons. Another substantial number of GHR-expressing neurons were distributed in the thalamus region. These observations are consistent with the expected expression pattern of GHR in the brain (8,12,13), and for the first time enabled us to study the function of specific GHR populations throughout the brain.

Evidence for the importance of GH-responsive neurons in the hypothalamus in modulating metabolism was reported in several studies (35-38). We have recently identified a unique population of nutrient-sensing leptin receptor (LepRb)-GHR expressing neurons that regulate hepatic glucose production and lipid metabolism, suggesting that these neurons are crucial for the metabolic functions of GHR-neurocircuitry (18). LepRb neurons co-express GHR in the ARC, DMH, and LHA, suggesting the role of GHR in these neurons as an integrating site of glucose metabolism regulation. In the ARC there is minimal overlap between AgRP neurons and SST neurons, with some transcriptional similarities between these neurons such as in their synaptic circuitry and function (31,39). Recent single-cell analysis of ARC neurons demonstrated that GHR is highly expressed in the tight cluster of AgRP^+^/SST^+^ neurons together with corticotropin-releasing factor receptor 1 (*Crhr1)* (31), suggesting the potential role of these ARC neurons in GH neurocircuitry.

The circuitries engaged by GHR^ARC^ neurons involve several neuroendocrine populations such as the SST, GHRH, and AgRP since GHR^ARC^ neurons are co-localized with these cells in the ARC. The majority of GHR^ARC+^ neurons in the ARC are also pSTAT5 immunoreactive after GH treatment, confirming their sensitivity to GH. We showed that chemogenetic activation of GHR^ARC+^ neurons modulated both glucose metabolism and energy homeostasis indicating that GHR^ARC+^ neurons lie within energy balance and glucoregulatory neurocircuits. While our current studies do not indicate which specific neuronal subpopulations within GHR^ARC^ are responsible for controlling each of these distinct physiological responses, genetic deletion of GHR in AgRP neurons did not affect glucose metabolism or energy homeostasis (19), indicating that the role of GHR in AgRP^-^ populations in the ARC is to coordinate these responses. GHR^ARC^ neurons only partially overlap with SST and GHRH neurons, thus the contribution of these ARC neuronal populations to GHR^ARC^-mediated glucose metabolism and energy homeostasis modulation remains to be clarified.

One of the established effects of GH in the ARC - is inhibition of its own secretion, as part of an auto-feedback circuit, involving the interrelationships between SST, GHRH, and AgRP/NPY - expressing neurons through GHR (40). This might be particularly important as properly regulated neural circuits within the GH axis modulate GH release under fed and fasting states (41), while the imbalance between these networks might be part of multiple maladaptive endocrine changes responsible for metabolic alterations in obesity. Our chemogenetic studies indicate that the hypothalamic GHR axis in ARC promotes glucoregulatory responses by enhancing glucose tolerance, and suggest that GHR^ARC^ neurons represent a distinct neuronal population within the GH axis that play a crucial role in the regulation of glucose metabolism. This effect is complementary to the counter-regulatory enhancing effect of GH axis during hypoglycemia (42).

GHR^ARC^ neurons represent a heterogeneous population, which includes neurochemically-defined neurons that control specific physiologic functions. For example, acute chemogenetic activation of AgRP neurons alters food intake and decreases energy expenditure (25). Additionally, activation of AgRP neurons acutely impairs systemic insulin sensitivity by inhibiting glucose uptake in brown adipose tissue (43). However, while the majority of GHR^+^ neurons in the ARC colocalize with AgRP neurons, *GHR* represents only a very small cluster within AgRP neuronal population (31); therefore, it remains possible that other GHR^+^ neuronal populations in the ARC contribute to GHR^ARC^-mediated modulation of energy balance. In support, chemogenetic activation of ARC-SST neurons, or intracerebroventricular (i.c.v.) infusion of SST analog acutely and significantly increases feeding responses (31,44). Infusion of SST analog also increases energy expenditure, drinking behavior, and lowers glycemic values (44), similar to the chemogenetic activation of GHR^ARC^ neurons. These findings indicate some functional similarities between SST and GHR^ARC^ neurons whose activation is sufficient in driving feeding, glucoregulation, and energy balance.

In summary, we have generated the GHR^tdTom^ mouse model to characterize the anatomical localization of brain-wide GHR expression. Using the GHR^tdTom^ mouse model we demonstrate that GHR^ARC^ comprises a unique neuronal population capable of controlling energy balance and glucose metabolism. While the significance of this ARC subpopulation of GH-responsive neurons in the control of certain aspects of energy balance and glucose regulation remains to be elucidated, our study emphasizes the role of GH axis as an essential hypothalamic center in regulating metabolic functions and provides a resource for studying the biology and functionality of GH-responsive neuronal populations in the brain.

## Abbreviations

CNS: central nervous system
GH: growth hormone
GHR: growth hormone receptor
STAT5: signal transducer and activator of transcription 5
POMC: proopiomelanocortin
ARC: arcuate nucleus of the hypothalamus
DMH: dorsomedial hypothalamic nucleus
LHA: lateral hypothalamus
PVH: paraventricular hypothalamic nucleus
SST: somatostatin
GHRH: growth hormone releasing hormone
AgRP: agouti-related peptide

## Author contributions

JBML, LKD, carried out the research and reviewed the manuscript. IA, OD, CU, MK, and, MK carried out the research. MS designed the study, analyzed the data, wrote the manuscript, and is responsible for the integrity of this work. All authors approved the final version of the manuscript.

## Acknowledgments

The authors thank Mohammed Alysofi and Hawraa Alromdhan for their technical assistance. This study was supported by the American Diabetes Association grant #1-lB-IDF-063, by a Feasibility Grant from the Michigan Diabetes Research Center (P30DK020572) and WSU funds for MS.

## Data Availability

The datasets generated during and/or analyzed during the current study are available from the corresponding author on reasonable request.

---

**SUPPLEMENTARY FIGURE 1.**
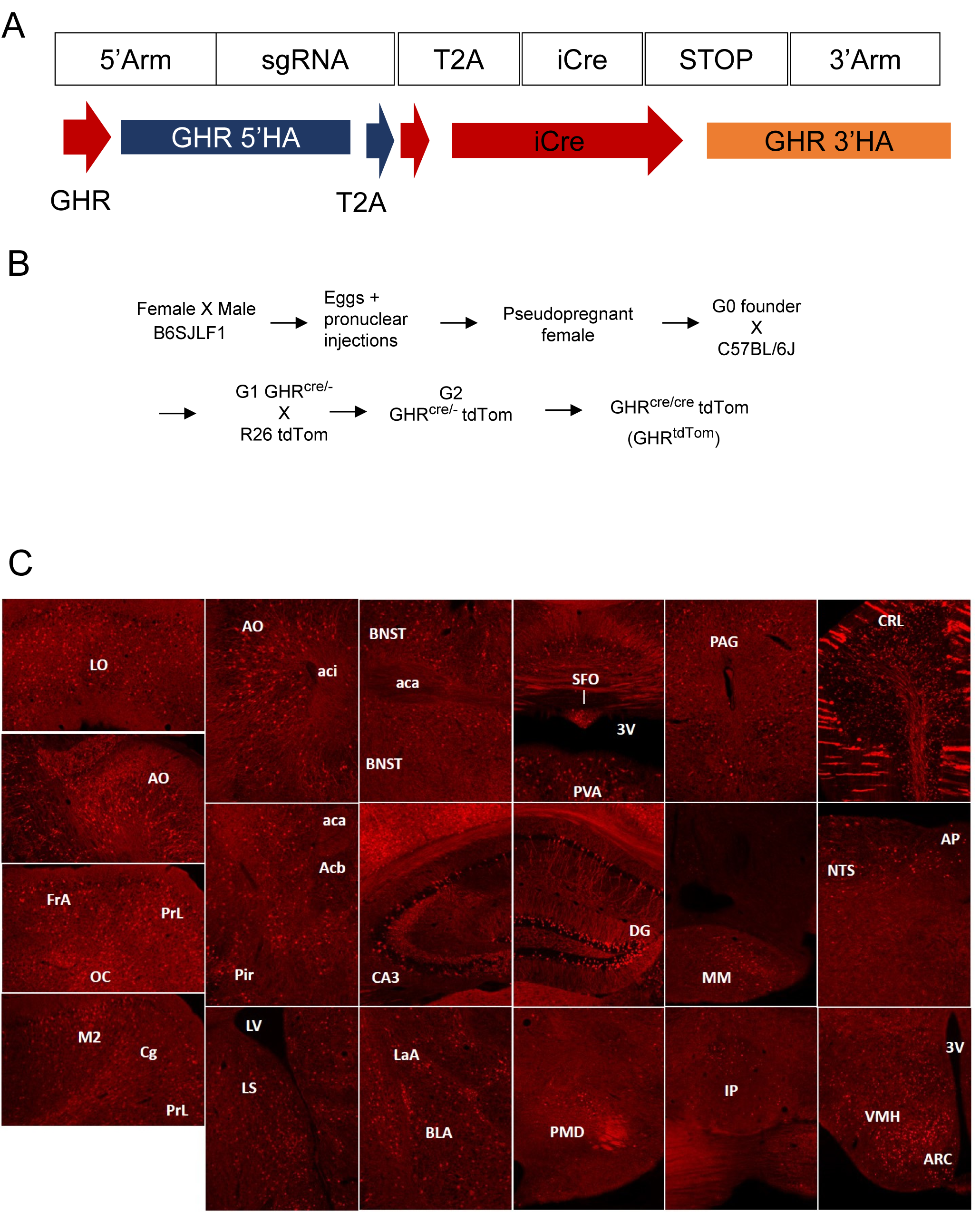

**SUPPLEMENTARY FIGURE 2.**
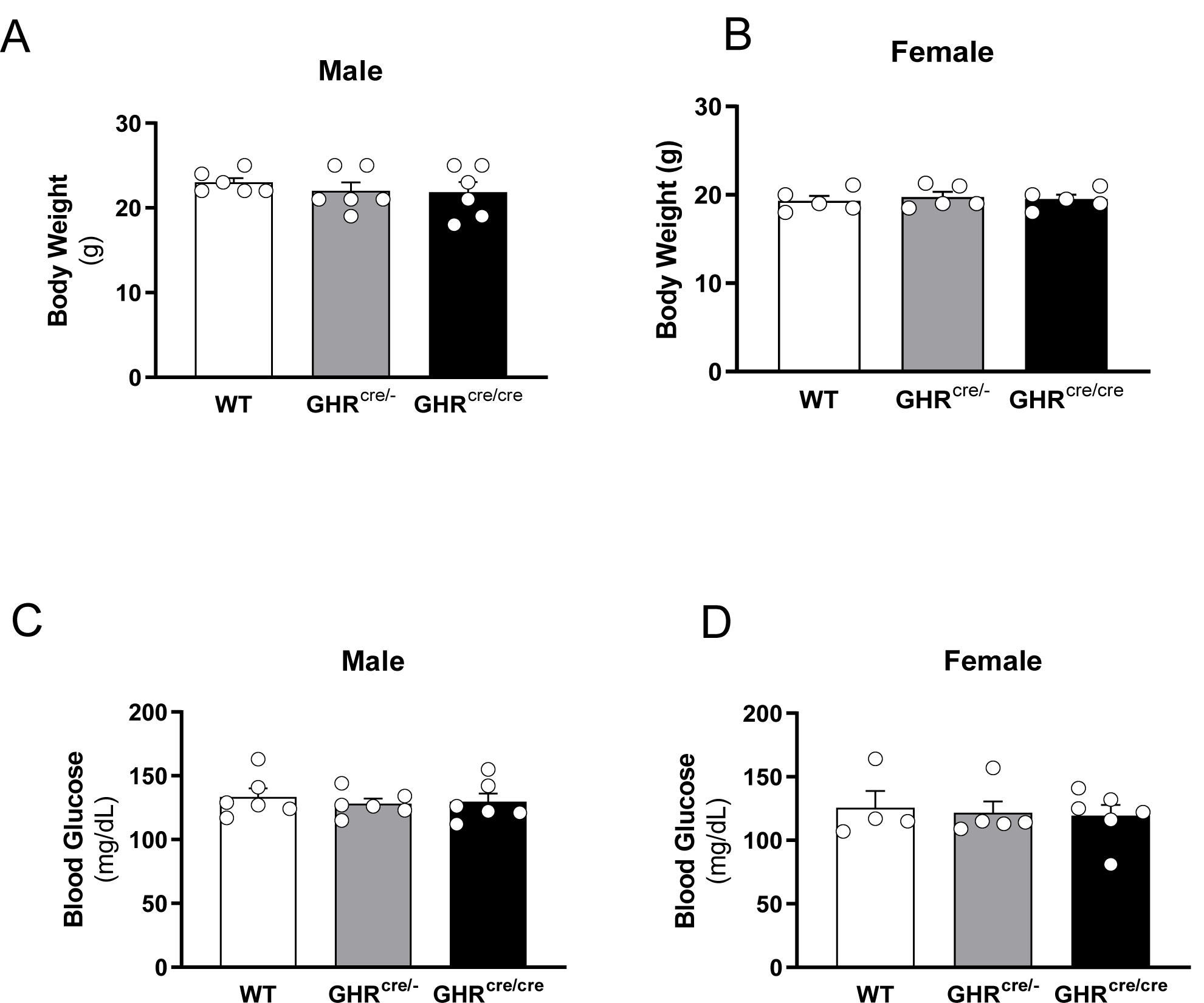

**SUPPLEMENTARY FIGURE 3.**
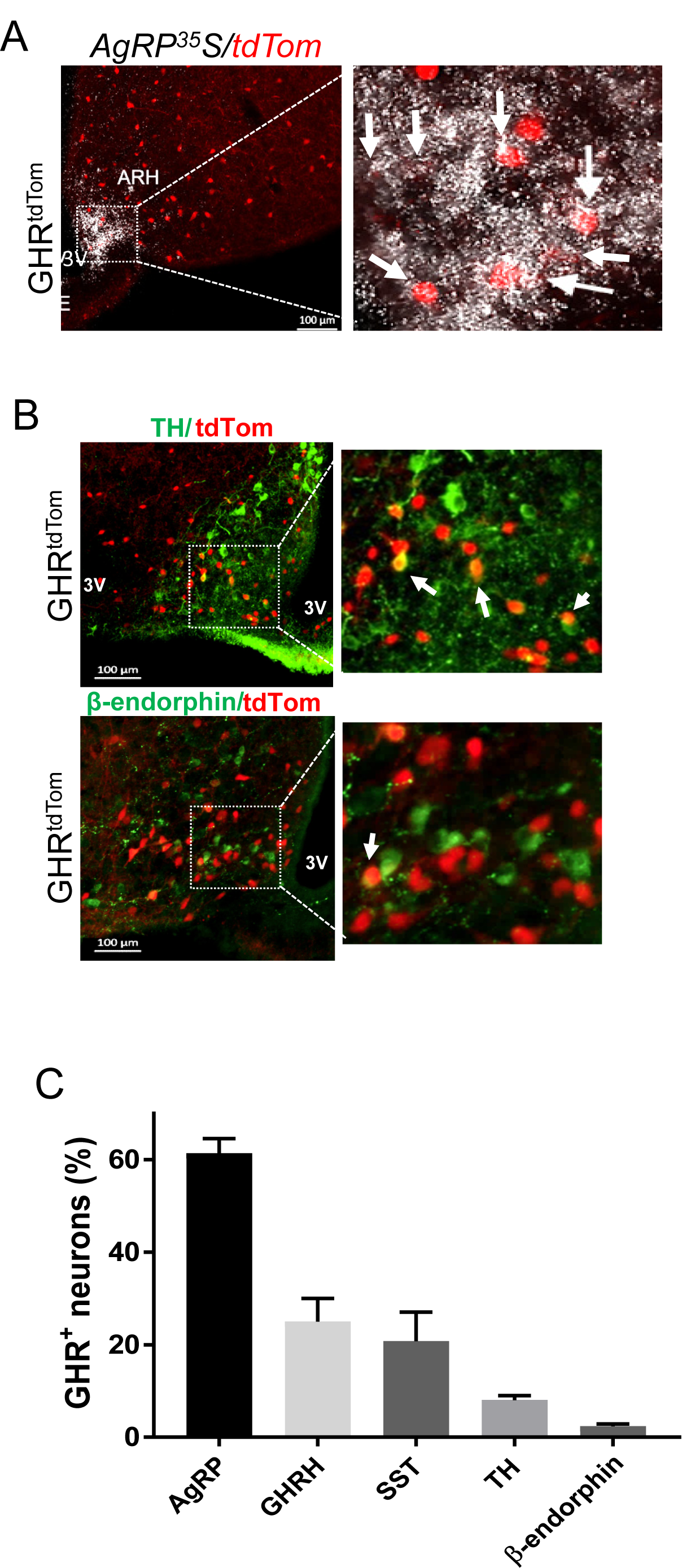

